# Patient Informed Domain Adaptation Improves Clinical Drug Response Prediction

**DOI:** 10.1101/2021.08.07.455527

**Authors:** Ioannis Anastopoulos, Lucas Seninge, Hongxu Ding, Joshua Stuart

## Abstract

In-silico modeling of patient clinical drug response (CDR) promises to revolutionize personalized cancer treatment. State-of-the-art CDR predictions are usually based on cancer cell line drug perturbation profiles. However, prediction performance is limited due to the inherent differences between cancer cell lines and primary tumors. In addition, current computational models generally do not leverage both chemical information of a drug and a gene expression profile of a patient during training, which could boost prediction performance. Here we develop a Patient Adapted with Chemical Embedding (PACE) dual convergence deep learning framework that a) integrates gene expression along with drug chemical structures, and b) is adapted in an unsupervised fashion by primary tumor gene expression. We show that PACE achieves better discrimination between sensitive and resistant patients compared to the state-of-the-art linear regularized method (9/12 VS 3/12 drugs with available clinical outcomes) and alternative methods.

## INTRODUCTION

Precision medicine promises to revolutionize cancer treatment by improving clinical drug response (CDR) prediction. CDR prediction could be greatly facilitated by cutting-edge high-throughput sequencing technologies, which provide comprehensive and individualized omics profiles. Based on these omics profiles, several CDR prediction approaches have been proposed. For instance, Tissue-Guided Lasso TG-LASSO^1^ integrates tissue-of-origin information with gene expression profiles for CDR prediction. DeepDR^2^, on the other hand, predicts CDR from mutation and expression profiles.

However, as another crucial component for CDR prediction, the chemical properties of drugs have been under-utilized. Although the traditional Morgan Fingerprint molecular representation^3^ has been used to integrate drug chemical information for CDR prediction, it cannot adaptively learn alternative representations of drug chemical properties as it is a static representation of the molecule and does not dynamically extract features for the desired prediction task. For example, CDRscan, similarly to TG-LASSO, does not take advantage of the diverse patient RNA-Seq profiles that are published on The Cancer Tumor Atlas (TCGA), and is not evaluated to address if the model can be applied for drug response prediction to patients, which is what such a model would be used for in practice. In addition, CDRscan uses Morgan Fingerprints to represent key molecular substructures using an explicitly defined featurization. A limitation of this specific methodology is its inability to adaptively learn alternative representations that may be beneficial to the particular task in hand^4^. DrugCell also uses Morgan Fingerprints to represent drugs along with an Visible Neural Network (VNN) embedded in Gene Ontology (GO) terms, which provides interpretable results^5^, but it is also not evaluated on patients.

Graph Convolutional Network (GCN)^6^ representations emerge as a powerful alternative for encoding drug chemical properties. GCN adaptively learns chemical information by generalizing the convolution operation from a grid of pixels to a graph, where each node can have a variable number of neighbors. GCNs have been used to explore drug-target interactions and side effect predictions - the two most important factors for developing a new drug. For instance, Decagon uses GCN to predict potential side effects of a drug ^7^. Such methodological advances provide novel insights in incorporating drug chemical properties during CDR prediction.

The majority of CDR prediction algorithms are trained with cancer cell line (CL) drug preturbation profiles. CLs have long served as models to study molecular mechanisms of cancer, because they maintain valuable molecular information of the primary tumor from which they were derived. CLs offer the advantages of being easily grown, relatively inexpensive, and amenable to high-throughput assays. Data generated from CLs can then be used to link cellular drug response to molecular features, where the ultimate goal is to build predictive signatures of patient outcomes ^8^. Various models have been developed to predict patient CDR from the molecular profiles of CLs ^9–11^. However, these models only show limited success in certain drugs ^12^ ^13^. Therefore, developing a model based on CL molecular features to predict CDR in patients for most drugs remains challenging ^14^. One major difficulty for such cross-domain CDR prediction is the prominent differences between cell lines and primary tumors ^15–19^. Recent advances in domain adaptation aim at aligning domains to tackle domain alignment problems, such as batch effect correction to reconcile differences across laboratories and studies ^20^. Mean Maximum Discrepancy (MMD) ^21^ has shown promising results in aligning domains in an unsupervised manner ^22^. Such a technique could be used to align CL and patient tumors in developing drug response models that are more clinically focused.

Inspired by the advanced GCN-based drug chemical information encoding, as well as the MMD-based domain adaptation, we develop a drug response predictor using Patient Adaptation and Chemical Embedding (PACE). This deep learning framework uses a GCN to dynamically learn chemical information of each compound, and is adapted to implicitly align the CL expression representation with that of a patient sample of the same tissue of origin. Thus, the model does not assume that CL and patient samples are drawn from the same distribution. We achieve this by using Mean Maximum Discrepancy (MMD) to align the latent spaces produced by CL expression and patient expression vectors. Such a technique has been successfully applied in previous studies ^23–25^ in transferring generalizable features across domains ^26^. We trained our model on 142,351 of CL-drug-IC_50_ (CDI) pairs where each CL vector was paired with a random patient expression vector of the same tissue of origin. We used a Graph Convolutional Network (GCN) to encode drug information and pair it with the patient adapted expression information in order to predict IC_50_. To evaluate our model, we collected a curated CDR dataset with patient outcome information on response to 12 drugs in total. Our model achieved superior performance (9/12 drugs) compared to alternative PACE models and the state-of-the-art linear method (3/12 drugs) in significantly discriminating between sensitive and resistant patients.

## RESULTS

### Overview of PACE and evaluation strategy

The PACE deep learning framework consists of three modules: an Expression Module (EM), Drug Module (DM), and Prediction Module (PM). As shown in Figure 1A, the EM is composed of fully connected layers and learns highly informative features from gene expression vectors. The DM is composed of a GCN and learns highly informative features for each atom from the graph representation of chemical compounds. Atom-level features are then aggregated to represent information about the compound as a whole (see METHODS). Given the success GCNs have had in computational chemistry and biology applications ^27–29^, we posited that the DM could learn a general graph embedding that would extend to drugs unseen during training. The PM is composed of a fully connected layer and takes the information learned from the EM and DM as input to predict log(IC_50_). We included gene expression information, the drug used, and the associated IC_50_ value from the Genomics of Drug Sensitivity in Cancer (GDSC) project ^30^. The model was trained with “CDI tuples” -- <u>C</u>ell line, <u>D</u>rug SMILES, <u>I</u>C_50_ -- indicating which drug was applied to a particular cell line and what that cell line’s response was to the drug.

**Figure 1.**
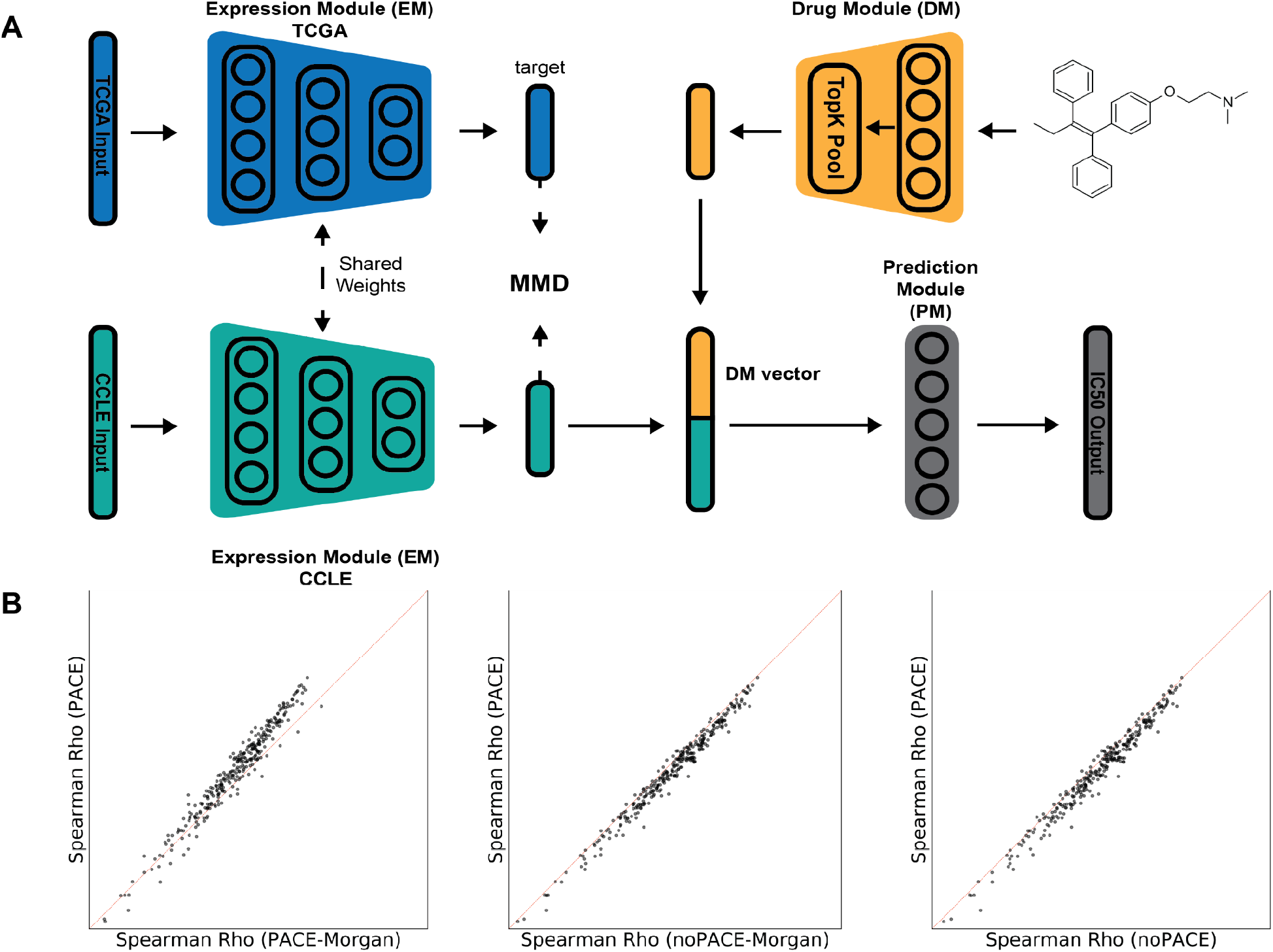
Deep learning model architecture. (A) Graphic overview of the proposed deep learning framework. Expression Module (EM) extracts highly informative features for the input expression vectors for both CCLE and TCGA via shared weights. These two compact expression representations are compared with each other via Mean Maximum Discrepancy (MMD) to diminish the distance between them, thereby aligning the two representations. The Drug Module (DM) encodes the molecule and pools the most informative nodes (atoms) to also create a highly informative compact representation. Finally, the CCLE expression representation and the drug representation are concatenated together and passed to the Prediction Module (PM) that makes the final log(IC_50_) prediction for each CDI pair. (B) Per drug predictive performance, which is quantified by theSpearman Rho between actual and predicted log(IC_50_) across all perturbed cell lines. GCN-MMD is compared against MORGAN-MMD (left), GCN-NoMMD (middle), MORGAN-NoMMD (right). Each point represents a drug. Points above the diagonal represent better performance by GCN-MMD.

Our goal is to extrapolate drug response from cell lines to patients. Hence, the model needs to generate an out-of-distribution (OOD) embedding space (for patient samples) representing a distribution not present in the training data (of cell lines). Inspired by ^23^, and recent advances in the field of domain adaptation^26^, we used maximum mean discrepancy (MMD) to adapt the latent distribution produced by the EM so that cell line gene expression is aligned to patient gene expression. Each CL was paired with a TCGA tumor sample’s gene expression vector of the same tissue of origin (see Supplemental Table 1/2), which has been shown to play a key role in a tumor’s treatment and progression. Restricting cell lines to match the tissue of origin of primary tumors resulted in 531 cell lines treated across 310 drugs, amounting for 142,351 CDI pairs. Cell lines and tumors that did not have a matching tissue of origin were not included in training. Each cell line was paired with a random primary tumor sample of the same tissue with the goal of creating a general enough adaptation of EM’s latent space.

To test the efficacy of adapting the EM with patient gene expression via MMD, we constructed a non-adapted version of PACE for comparison purposes. In addition, we also compared our model to one in which the DM uses Morgan Fingerprints (MorganFP), representing a more conventional molecular encoding. Altogether, we created three alternative models closely related to PACE -- PACE-Morgan, noPACE, and noPACE-Morgan (see Table 1). Alternative models with an adapted EM should provide a poorer fit to the cell line data, yielding poorer performance in cell lines, compared to non-adapted models because the adapted models attempt to fit the distribution of both cell lines and patients. On the other hand, the adapted models should perform better in the patient setting.

**Table 1.** All alternative models used and their specifications.

| EM | DM | Name |
| --- | --- | --- |
| Adapted | GCN | PACE |
| Adapted | MorganFP | PACE-Morgan |
| Not adapted | GCN | noPACE |
| Not adapted | MorganFP | noPACE-Morgan |

### Combining domain adaptation and graph encoding preserves performance in cell lines while increasing the accuracy of predicting patient response for more drugs

We compared the Spearman Rho achieved by our proposed PACE model to all the other variations for cell lines treated by each drug in our dataset (Figure 1B). We found that although PACE does better compared to PACE-Morgan for the majority of the drugs (points above the diagonal), the non adapted versions (noPACE and noPACE-Morgan) achieve a higher correlation (points below the diagonal). Nevertheless, when comparing across all 142,351 CDI pairs, the proposed model and the alternatives achieved comparable results (Supplementary Fig. 1). This suggests that MMD adaptation preserves the prediction performance of drug response in CL while yielding superior discrimination performance in the patient setting (shown next).

In addition to the models described in the previous section, we compared PACE to the state-of-the-art TG-LASSO^1^ model, which is a linear regularized method to predict *clinical drug response* (CDR). To evaluate all of the models in the patient setting, we followed the same evaluation presented in the TG-LASSO study^1^. We used the same curated CDR dataset consisting of 531 patients treated across 24 drugs labeled with the type of response indicated for each patient. The majority of patients in this dataset (70%) were treated with a single drug, while the rest were given two or more. Patients with stable disease and clinical progressive disease were labeled as resistant (R), whereas those with partial or complete response were labeled as sensitive (S). After following the same filtering steps and retaining samples for which we had expression information, 506 patients across 12 drugs remained. To measure the performance of the methods, we asked if their predicted log(IC_50_) drug response (a continuous measure) correlated to the drug response labels (R/S) (a categorical measure) in the CDR dataset (see METHODS). Specifically, a one sided Mann-Whitney U test was used to determine whether the predicted log(IC_50_) for the true resistant (R) patients is significantly larger than that of true sensitive (S) patients.

As shown in Figure 2A, MMD adaptation produces an embedding that can discriminate between resistant and sensitive patients across more drugs compared to all other models that lack such adaptation. The combination of patient information adaptation with MMD and GCN for drug embedding had better correlation to patient response than all the alternative methods examined (Figure 2A, Supplementary Fig 2). Specifically, PACE showed significant discrimination between resistant and sensitive patients (p<0.05) for nine out of the twelve drugs compared to six by noPACE (Figure 2A). Similarly, PACE-Morgan predicted six drugs, compared to five by noPACE-Morgan (Supplementary Fig. 2). TG-LASSO was the worst performing method with three drugs predicted significantly.

**Figure 2.**
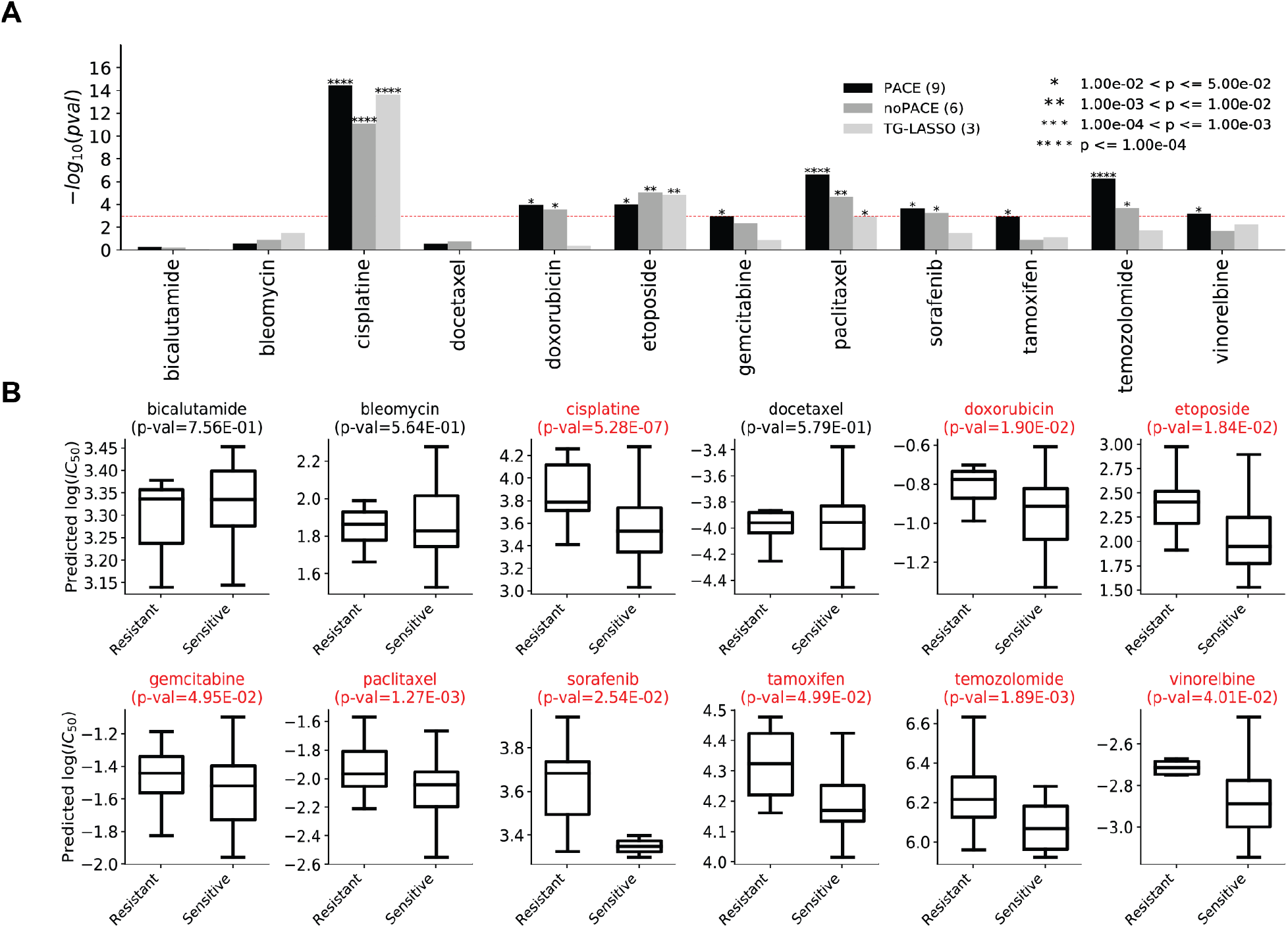
TCGA CDR performance comparison with framework variants and with TG-LASSO. We compare our model with the unregularized alternative and with the state-of-the-art TG-LASSO linear method. GCN-MMD yields the best resistant-sensitive discrimination performance in the TCGA CDR dataset (A) Bar plots showcasing p-value, corresponding to the one-sided Mann-Whitney U test, determined by averaging 10 independent predictions made by each model. (B) Box plots reflecting the distribution of estimated log(IC_50_) values using GCN-MMD for resistant or sensitive patients. The p-values here also correspond to a one-sided Mann-Whitney U test.

We also observed that regardless of the method used, cisplatin, etoposide, and paclitaxel were predicted correctly. In contrast, bicalutamide, bleomycin, and docetaxel were not predicted correctly by any of the methods. We further observe that CDR prediction using MMD adaptation improved CDR prediction for cisplatin, doxorubicin, gemcitabine, paclitaxel, tamoxifen, temozolomide, and vinorelbine suggesting that tissue of origin may play a crucial role for these drugs. However, for gemcitabine, tamoxifen and vinorelbine, using the MorganFP drug encoding did not achieve significant CDR prediction in patients, even with MMD adaptation, further suggesting that the appropriate drug embedding is needed for such a task (Supplementary Fig. 2).

Lastly, we found that for the hard to predict bicalutamide PACE and PACE-Morgan produced predicted IC_50_ values with the correct direction (predicted IC_50_ for resistant patients should be higher than that of sensitive patients) (Figure 2B, Supplementary Figure 3A). The non-adapted variants of PACE (noPACE, noPACE-Morgan) produced predicted IC_50_ values with the incorrect direction (Supplementary Figure 3B/C). This is also evident by the direction of the difference between the median of sensitive predicted IC_50_ and sensitive predicted IC_50_, which we term as *ΔIC*_*50*_ (Supplementary Table 3). The *ΔIC*_*50*_ for bleomycin was also observed to be the most negative in the PACE models compared to the noPACE models. PACE-Morgan produced the most negative *ΔIC*_*50*_ for docetaxel compared to the noPACE models, however for this hard to predict drug PACE produced a *ΔIC*_*50*_ in the wrong direction.

Taken together, these results suggest that the combination of patient adaptation via MMD and a combination of the chemical embedding learned from GCN produced a highly informative model that can be extended to the patient setting.

### Cell line diversity is more important than drug diversity for patient CDR prediction

Next, we asked if gene expression information or drug information has a bigger impact in predicting drug response in patients. To this end, we created two different drop out experiments -- one where all the CDI pairs for a *cell line* were withheld and another in which all the CDI pairs for a *drug* were withheld. For the cell line dropout experiment, we created training sets with 20%, 40%, 60%, and 80% of the total cell lines (531). For the drug dropout experiment, we measured the performance of a method in predicting the twelve CDR drugs having never seen those same drugs during training. To this end, we removed the 12 drugs present in the CDR dataset and then created training sets in cell lines that included 20%, 40%, 60%, and 80% of the remaining 298 drugs. For each of the training sets, the model was trained ten independent times on each fold. The ten independent cell line-trained models were then applied to the patient CDR dataset, and the average predicted log(IC_50_) was computed. The Mann-Whitney U test was used to evaluate the discrimination between labeled resistant and sensitive patients (see METHODS).

As summarized in Figure 3A, lack of gene expression information had a bigger impact compared to lack of drug information across all drugs in our CDR dataset. This is likely caused by the vast difference in complexity and variance between the gene expression profiles and the compound structures. The robustness (measured by the variance of the p-value across 10 fold cross validation) of the model suffers more with 20% of the CLs included in training compared to the same percentage of drugs included in training (Supplementary Figure 4A/B). Addition of more CLs in the training set drastically improves robustness of the model as shown by the decreasing variance of the p-value across all 10 folds, indicative of the crucial role expression information plays in predicting drug response (Supplementary Figure 4A). This result also suggests that the GCN needs a small amount of graph examples in the training set to be able to generalize well to new graphs not seen in the training set. Additional graph examples improved the robustness of the prediction of all but five drugs: bicalutamide, bleomycin, gemcitabine, sorafenib, and tamoxifen in the CDR dataset (Supplementary Figure 4B) with the variance in p-value decreasing slightly. We conducted an additional experiment, where only the 12 CDR drugs were removed from training and reported minimal reduction in CDR prediction performance (Figure 3B). It is crucial to note that the small improvements in CDR prediction from additional training molecules can be explained by the fact that most of the CL have been treated by most of the drugs (median number of CL = 498.5). This means that most of the variability in gene expression has been seen by the model even when 20% of the drugs are included in training (Supplementary Figure 4C), which leads to robust results in the CDR dataset and is consistent with what we observed in the CL dropout experiment.

**Figure 3.**
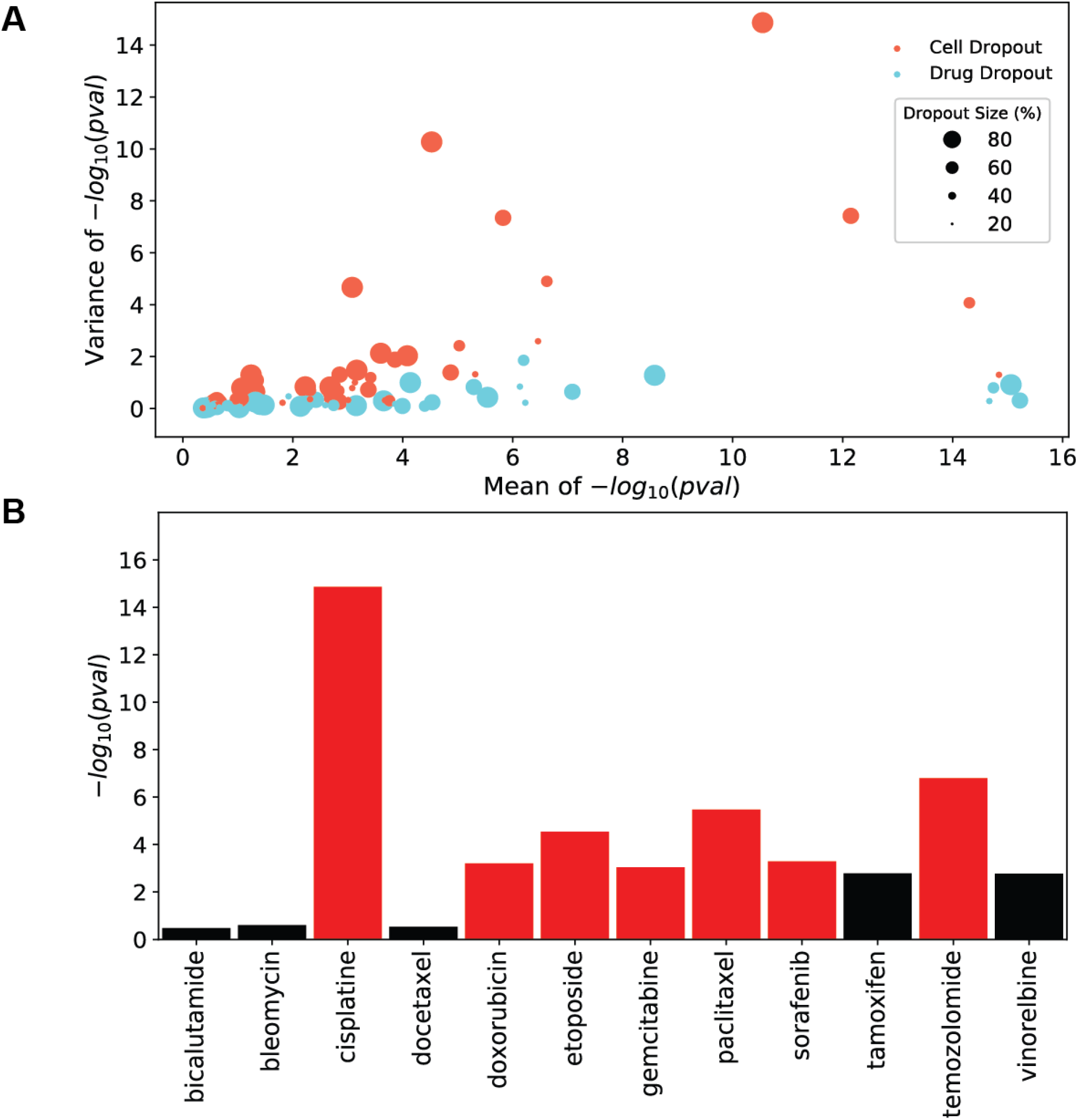
Drug/Cell Line 10 fold cross validation dropout experiment. Dropping data, drug-wise and CL-wise to test the limits of the model’s OOD inference ability in a 10-fold cross validation fashion. For each fold the training was repeated 10 independent times. (A) Showing the variance of the −log(p-val) determined by the Mann-Whitney U test for the difference between the predicted log(IC_50_) between resistant and sensitive patients across 4 conditions: 20% of CL retained in training, 40% of CL retained in training, 60% of CL retained in training, and 80% of CL retained in training. (B) Dropping only the 12 drugs in the CDR dataset. Variance results are displayed for all 12 drugs in the CDR dataset.

### Top Predictions Recapitulate Knowledge on Targeted Therapy

Most of the drugs with CDR in patients that were tested here can be classified as chemotherapy agents, with the exception of sorafenib (VEGFR inhibitor) and tamoxifen (*ESR1* inhibitor). To assess the performance on targeted agents for well characterized cohorts, we carried out an *in-silico* analysis on drugs with known biomarkers of response. We used mutations as biomarkers of response for the TCGA cohorts where expression and mutation information were available. For the cohorts where these were not available we used expression as a biomarker of response to confirm that the model learns biologically meaningful information. The idea here is that as a target gene’s expression increases, the drug’s predicted IC_50_ should decrease accordingly, indicating an increase in sensitivity.

We collected mutation information from TCGA breast cancer (BRCA), melanoma (SKCM) samples and LUSC/LUAD cohorts (combined and abbreviated as LUNG). We used mutation information as a biomarker of sensitivity. We tested trametinib, olaparib, dabrafenib and gefitinib on all of the aforementioned cohorts. Trametinib is a MEK inhibitor and used to treat SKCM. Olaparib is a PARP1 inhibitor and used to treat *BRCA1*- or *BRCA2*-mutated breast and ovarian (OV) cancers. Next, we examined the correlation between a drug’s predicted log(IC_50_) and its target gene’s expression (after Z-score transformation of the gene expression values). We specifically examined OV in this way due to the fact that we could not collect sufficient OV samples with predicted *BRCA1* or *BRCA2* mutations, and thus could not use *BRCA1/2* mutation as a biomarker for OV and olaparib.

As expected, *BRCA1* mutant samples were predicted to be more significantly sensitive to olaparib compared to *BRCA1* WT samples (Figure 4A), *NRAS* and *MAP2K1* SKCM mutants were predicted to be significantly more sensitive to trametinib compared to SKCM WT samples (Figure 4B/E). *NRAS* SKCM mutants were additionally predicted to be more sensitive to dabrafenib compared to *NRAS* WT samples (Figure 4C). Olaparib has been previously shown to be effective in *ATM* mutated BRCA patients. Although our model was not able to predict this association correctly, SKCM *ATM* mutated samples were predicted to be sensitive to olaparib (Figure 4D). Lastly, LUNG cancer *NRAS* mutated samples were predicted to be more sensitive to dabrafenib (Figure 4F).

**Figure 4.**
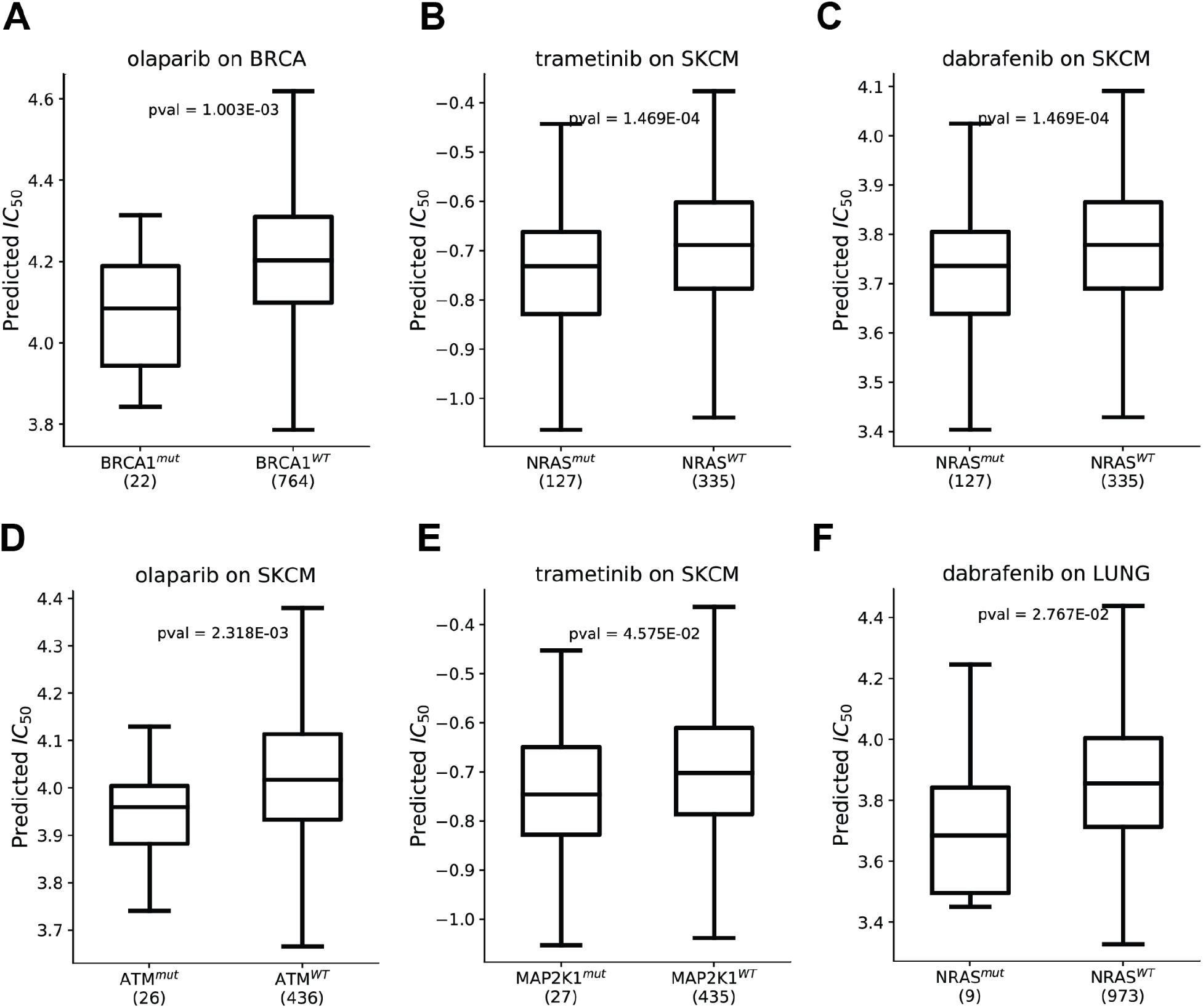
Functional Analysis. (A) Predicted log(IC50) for BRCA1 mutant and wild-type (WT) breast cancer (BRCA) samples in-silico treated with olaparib. P-value corresponds to the one-sided Mann Whitney U test discriminating between mutant and WT predicted log(IC50). (B) Predicted log(IC50) for NRAS mutant and wild-type (WT) melanoma (SKCM) samples in-silico treated with trametinib. (C) Predicted log(IC50) for NRAS mutant and wild-type (WT) melanoma (SKCM) samples in-silico treated with dabrafenib. (D) Predicted log(IC50) for ATM mutant and wild-type (WT) melanoma (SKCM) samples in-silico treated with olaparib. (E) Predicted log(IC50) for MAP2K1 mutant and wild-type (WT) melanoma (SKCM) samples in-silico treated with trametinib. (F) Predicted log(IC50) for NRAS mutant and wild-type (WT) LUSC/LUAD (LUNG) samples in-silico treated with dabrafenib.

Studies have pointed to *PARP* expression as a promising biomarker of olaparib response^31^. When we examined the correlation between *PARP1* z-score and the predicted log(IC_50_) per disease, OV had a significantly negative Spearman Rho (rho=−0.48) (Supplementary Figure 5A). Testicular cancer (TGCT) showed the strongest negative correlation between *PARP1* expression and predicted IC_50_ for olaparib, which has recently been in clinical trials in combination with chemotherapy for TGCT^32^.

We further examined the predicted response of lapatinib and the correlation with the expression of its target genes, *EGFR* and *ERBB2* (gene expression first transformed. *In-vitro* studies have previously shown that lapatinib inhibits cell proliferation and migration of breast cancer cell lines expressing different levels of *EGFR* and *ERBB2*, and that cells overexpressing *ERBB2* were more sensitive^33^. Interestingly, our model predicted *EGFR* expression as a stronger biomarker (Supplementary Fig. 5B) in BRCA patients compared to *ERBB2* expression (Supplementary Fig. 5C). Taken together, these results suggest that our model can recapitulate the relationship of well characterized drugs with the appropriate biomarkers, and their applicability in equally well-characterized cohorts.

## DISCUSSION

In this study, we presented a new deep learning framework that uses both a graph convolutional network (GCN) as a general encoding for drug information together with patient information to aid in out-of-dataset prediction. During training, the method aligns cell-line and patient gene expression domains using implicit tissue-driven adaptation together with drug information to derive highly informative features for drug response prediction.

We showed that adapting tumor information with maximum mean discrepancy (MMD) preserves performance in cell lines while improving the prediction of clinical drug response (CDR) in patients regardless of the drug encoding used. We found that GCN’s embedding extends to drugs that have not been seen in training. These results suggest that a combination of implicit tissue-driven adaptation and a highly flexible drug encoding lead to improved prediction performance in patient samples. Interestingly, we note that the drug dropout experiments revealed that only 20% (298) of drugs are needed to yield robust generalization performance. On the other hand, the cell line dropout experiments showed that a lack of cell line diversity during training greatly impacts generalization of drug response in patients. We further examined if our model can recapitulate some of the well known therapeutics for melanoma, breast cancer, and lung cancer. We found that the model was able to predict *MEK2* mutant melanomas as significantly more sensitive to trametinib, a MEK inhibitor, compared to the WT cohort. Similarly, *BRCA1* mutants in breast cancer were significantly more sensitive to olaparib, a first-line treatment to patients with such a mutation, compared to the WT cohort. #

Ideally, the type of models studied here should be trained on patient samples rather than surrogates such as cell lines. However, at this time, an adequate amount of patient data is lacking for any particular drug of interest as most patients receive the standard of care based on the tissue of origin. For example, we attribute the poor performance in predicting sensitivity to bicalutamide, bleomycin, and docetaxel to lack of adequate training data (Figure 2A). In the coming years, single cell sequencing should improve the performance of predictive models. For example, promising results have been published by the MIX-seq study in which the sequencing of cell lines before and after drug treatment has detailed the heterogeneity in response across individual cancer cells^36^. Together with single-cell sequencing, human-derived xenografts and 3D human organoids should complement cell line studies to add needed realism for classifier training; e.g. by including contributions from the microenvironment. #

The framework we presented can be extended to incorporate additional diverse biological data as it becomes available. As expected, we found that accuracy depended heavily on the presence of an appropriate cell line of a matching tissue type in the training data. Beyond extending the training data to cover more cell lines, which will increase the diversity of patients to which the method can be applied, other data types may also provide a boost in performance. For example, the current work focuses on gene expression and does not consider genomic alterations, such as mutations and structural variants, and the vulnerabilities that these may introduce. Genetic dependency data generated from the ACHILLES project ^37^ for example are now available for many of the same cell lines that our model was trained on. In theory, incorporating synthetic lethality prediction into the model should improve drug response prediction, as drugs that target synthetically lethal pairs should have a substantial impact on drug response. Incorporating protein level information could also lead to improvements in performance as many of the drugs target specific proteins whose expression may or may not be correlated with the gene’s RNA. The ongoing CPTAC project ^38^ is systematically quantifying protein levels and phosphorylation states in cancer patients from TCGA. In addition, it has been shown that proteome-level characterization of cell lines can aid in drug response prediction ^39^. It is therefore evident that addition of proteomic data to our model could have a significant impact on the prediction of drug response.

Lastly, increasing the interpretability of our model would be of great value. It would be very informative to developers of new drugs if they could predict the pathways affected by administration of a new treatment. Recent advances in developing more interpretable biological models^40,41^ should help models like ours in providing generalizable and interpretable results. Lastly, the GCN of our model uses only atom features for drug encoding. Other types of GCN, such as GINEConv from this study ^42^, are more expressive and use both atom and bond features, which could potentially create an even more generalizable drug embedding. We leave the exploration of the most appropriate GCN for this task and the inclusion of an interpretable EM to future studies.

## METHODS

### Overall Framework

Our model is an adapted dual convergence architecture that integrates gene expression information with drug structure aimed at generalizing clinical drug response (CDR) prediction in patients. It consists of three modules: Expression Module (EM), Drug Module (DM) and Prediction Module(PM). Highly informative representations of gene expression and drug structure are generated by the EM and DM, respectively. These representations are jointly passed to the PM where the log(IC_50_) prediction is made. The model takes as input a cell line (CL) expression vector (***x***_***c***_), a primary tumor expression vector (***x***_***t***_), and the compound that was applied on the CL. The way the compound is presented as input to the model is explained in the ***Morgan Fingerprint (MorganFP) Representation of Drugs*** and ***Graph Representation of Drugs*** sections.

### Expression Module (EM)

The EM consists of 2 fully connected layers of 1024, and 100 nodes with Rectified Linear Unit (ReLU) activation. BatchNormalization and Dropout of 0.35 are applied on each layer. During training, the EM produces latent representations for both CL and primary tumors via weight sharing as follows:

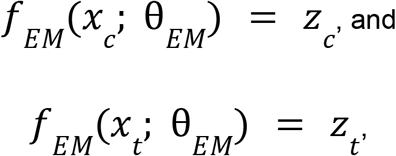

Where *Z*_*c*_ and *Z*_*t*_ represent the latent vectors of CL and primary tumor, respectively.

Inspired by the field of domain adaptation, and driven by the need to generalize drug response prediction to patients, we used a domain alignment method called Mean Maximum Discrepancy (MMD) ^15^. Specifically, the model tries to align *Z*_*c*_ to *Z*_*t*_ with the goal of making the cell line latent space more similar to the primary tumor latent space by minimizing the following loss:

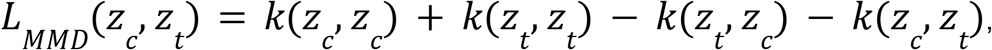

where*K*denotes the universal Gaussian kernel. Each *Z*_*c*_ and *Z*_*t*_ represent the same tissue-of-origin during training. Thereby, the model implicitly aligns CL and primary tumors in a tissue driven manner.

### MorganFP Representation of Drugs

We used the python library RDKit to generate Simplified Molecular Input Line Entry System (SMILES) strings, which describe the structure of a molecule using a single line of text, and compute MorganFP for each molecule in our datasets ^43^. SMILES strings are simple string annotations that describe the structure of the molecule. MorganFP is part of the Extended-Connectivity Fingerprints (ECFPs) family and are generated using the Morgan algorithm ^44,45^. These fingerprints represent molecular structures and the presence of substructures by means of circular atom neighborhoods (bond radius). In this study we used radius 2 and constructed a 2048 long bit vector for each molecule. A radius of 2 takes into account neighbors up to two atoms away when constructing the bit vector (fingerprint) of the molecule.

### Graph Representation of Drugs

We used RDKit to generate SMILES strings for each drug. Next, we represented the SMILES string for each compound {*c*_*j*_} ∈ *C* as a graph *G* = {*V*, *X*}, where *V* = {*ν*_*j*_} represents the set of nodes (nodes here are atoms on the molecule). An adjacency matrix *A* represents the topological structure of each molecule with *A*_*i,j*_ = 1 denoting a bond between two atoms, otherwise *A*_*ij*_ = 0. *x*_*i*_ ∈ *X* indicates the vector of features for each atom *ν*_*i*_ on the compound. The features (189 in total) used for each compound can be found in Table 2.

**Table 2.**
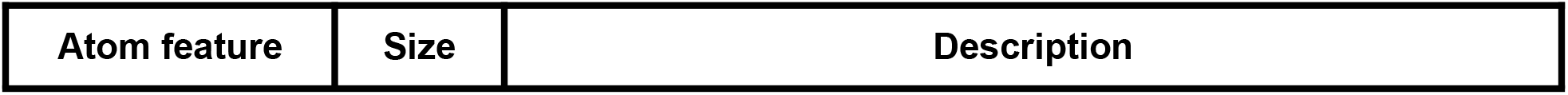
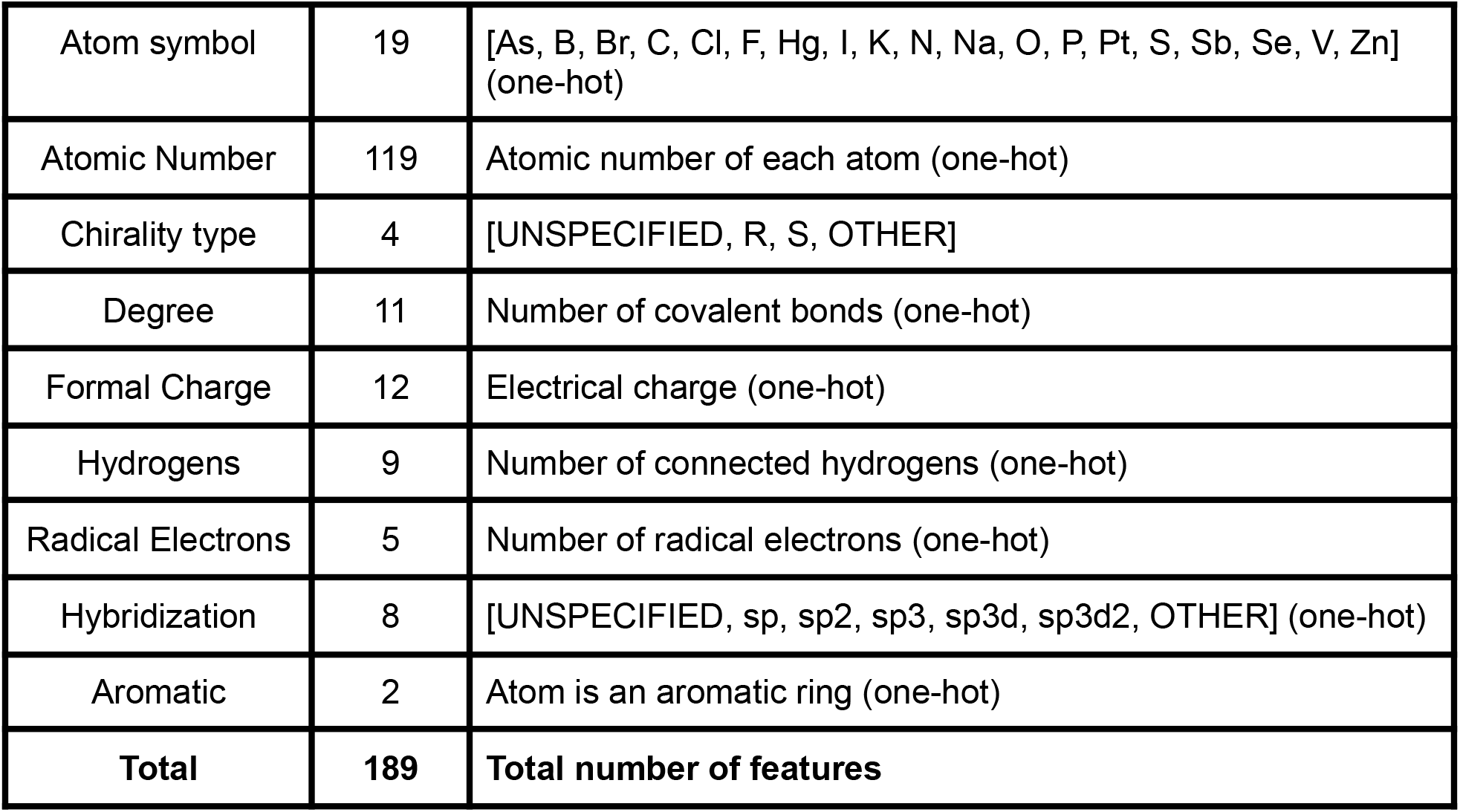
Description of Atomic Features.

| Atom feature | Size | Description |
| --- | --- | --- |
| Atom symbol | 19 | [As, B, Br, C, Cl, F, Hg, I, K, N, Na, O, P, Pt, S, Sb, Se, V, Zn] (one-hot) |
| Atomic Number | 119 | Atomic number of each atom (one-hot) |
| Chirality type | 4 | [UNSPECIFIED, R, S, OTHER] |
| Degree | 11 | Number of covalent bonds (one-hot) |
| Formal Charge | 12 | Electrical charge (one-hot) |
| Hydrogens | 9 | Number of connected hydrogens (one-hot) |
| Radical Electrons | 5 | Number of radical electrons (one-hot) |
| Hybridization | 8 | [UNSPECIFIED, sp, sp2, sp3, sp3d, sp3d2, OTHER] (one-hot) |
| Aromatic | 2 | Atom is an aromatic ring (one-hot) |
| **Total** | **189** | **Total number of features** |

### Drug Module (DM)

The DM of the model aims at extracting highly informative features from each molecule. This is done via either the MorganFP representation of the molecule, or the graph representation of the molecule. For the former, the DM consists of one fully connected layer, ReLU, BatchNormalization and Dropout. For the latter, we used the python library PyTorch Geometric to produce data-driven molecular features using GCN ^46^. In particular, we used the GCN architecture from ^47^. That architecture learns substructures of a given graph, and relationships between graphs, which is crucial in this study as we aim to generate a general embedding space for structurally diverse molecules presented in the drug response dataset. This type of GCN falls under the spatial GCN category, which can generalize the learned embedding to heterogeneous graphs ^48^,. We used one layer, followed by a pooling layer, which aggregates highly informative nodes on the molecular graph ^49^. The DM consists of one layer due to the small average size of the molecules (34 nodes). ReLU, BatchNormalization and Dropout were applied here as well.

The purpose of the GCN is to map each *ν*_*i*_ ∈ *V* to low dimensional vectors *Z*_*i*_ ∈ *R*. The formal mapping is as follows:

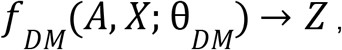

with *Z* ∈ *R*^*n×d*^ for compound *c*_*j*_, where *n* is the number of atoms, and *d* is the dimension of the latent space produced by the GCN.

Furthermore, to obtain a latent representation *Z*_*d*_ for graph *c*_*j*_, we computed both average and maximal features across *Z* and concatenated them with the following operation:

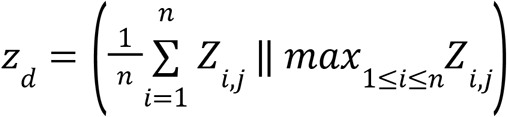

where *Z*_*d*_ ∈ *R*^*n×2d*^ and∥denotes the concatenation operator. The dimensionality is doubled due to concatenation of both average and maximal features for each graph.

### Prediction Module (PM)

The PM of the model consists of one fully connected layer, and aims at predicting log(IC_50_) using highly informative features derived from the EM and DM. As such, the operation carried out by the PM is the following:

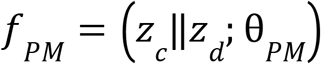

Our model updates the weights of EM, DM, and PM by minimizing the mean squared error (MSE),

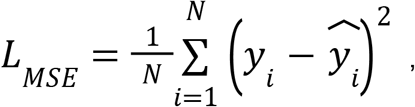

where *N* is the number of samples, between the observed and predicted log(IC_50_), denoted by *Y*and 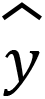, respectively, and *L*_*MMD*_.

Hence, the overall loss minimized by the model is:

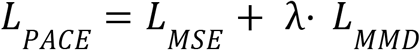

where λcontrols the tradeoff between the goals of aligning the CL latent space with the primary tumor latent space, and achieving an accurate predicted log(IC_50_).

### Training Procedure and Tuning

Our model was implemented in Python with the PyTorch API^50^ using the Adam optimizer ^51^ for gradient descent optimization. The training was allowed to proceed for a maximum of 200 epochs. To control for overfitting EarlyStopping was used to monitor the training loss for overfitting. Training was terminated after 10 epochs if the training loss was not further minimized after 10 consecutive epochs, with a delta of 0.05. Dropout was applied on a random 35% of nodes to further prevent overfitting. We used the Adam^51^ optimizer for gradient descent optimization with a learning rate of 1E-4. Given the stochasticity of the training procedure, and that we wanted to achieve considerable robustness with our model when predicting CDR of patients, we repeated the training 10 independent times.

Due to the computational expense of training, the number of layers for the DM and PM were fixed to one, and the number of layers of the EM were fixed to 2. We experimented with theλ, and with the number of drug nodes for the DM. We found that λ = 0. 01and 200 drug nodes were the best parameters for distinguishing sensitive from resistant patients in the CDR dataset

### CDR prediction in TCGA patients

We obtained the clinical drug response (CDR) of 531 TCGA patients across 24 drugs from this study ^17^. Following the same filtering steps as Huang et al. resulted in 12 drugs. Finally, after filtering for patients for which we had gene expression information resulted in 506 patients. Patients with “clinical progressive disease” or “stable disease” were labeled as resistant (**R**). Those with “partial response” or “complete response” were labeled sensitive (**S**). These are categorical variables, whereas our model predicts log(IC_50_) which is a continuous variable. To test how well our model can be extended to OOD samples, we grouped the predicted log(IC_50_) of each patient in the corresponding R or S bin. Then, we tested if the predicted log(IC_50_) of the R patients was significantly larger than that of the S patients by performing a one-sided nonparametric Mann Whitney U test. A summary of the number of R and S patients for each drug is shown in Table 3.

**Table 3.**
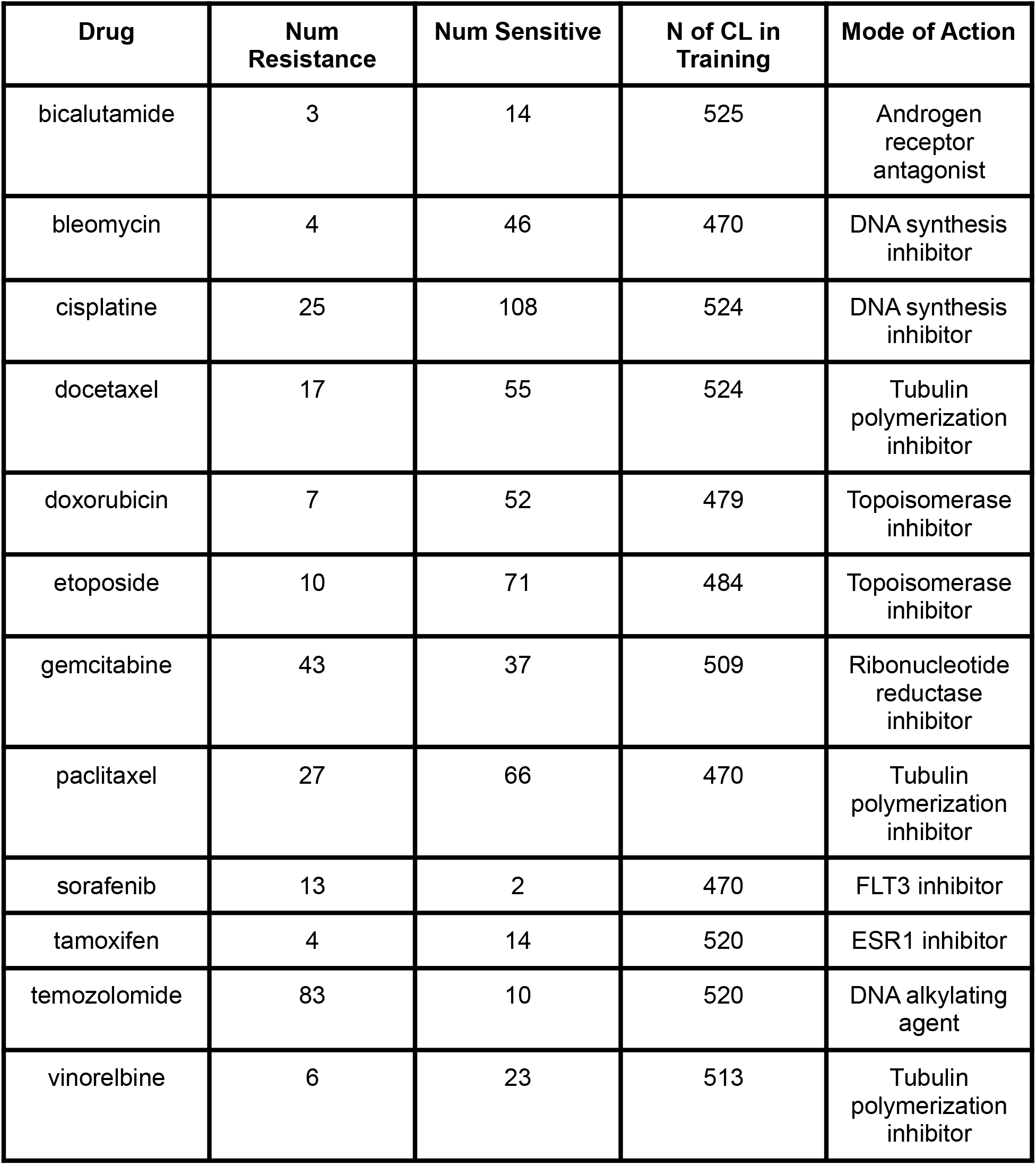
Number of Resistant and Sensitive Patients in TCGA CDR dataset.

### Drug/CL exclusion experiment

For dropout analysis, we created random train splits in a 10-fold cross validation. After training on each fold 10 independent times, we tested the generalizability potential of our model in the CDR dataset for each fold, thereby producing 10 p-values (see ***CDR prediction in TCGA patients)***. For drug-centered dropout analysis, we created train sets by first removing all 12 CDR drugs (bicalutamide, bleomycin, cisplatine, docetaxel, doxorubicin, etoposide, gemcitabine, paclitaxel, sorafenib, tamoxifen, temozolomide, and vinorelbine), and then retaining random 20%, 40%, 60%, and 80% of the remaining 298 drugs (310 drugs in total). Similar to the drug-centered dropout analysis, the CL-centered dropout was carried out in a similar manner without removing the 12 CDR drugs.

### Expression Datasets

We downloaded gene expression data of 1376 cell lines of the Cancer Cell Line Encyclopedia (CCLE) project, along with their metadata ^52^, and 10,536 TCGA pan-cancer tumors from the DepMap project ^53^ and UCSC Xena browser ^54^, respectively. All expression values were represented as log_2_(TPM+1), where TPM denoted transcripts per million reads of each gene in each sample. The gene space was intersected resulting in 31,501 common genes.

### Drug Response Datasets

We downloaded release 8.1 of the GDSC project containing drug response measured by the half maximal inhibitory concentration (IC_50_) from the DepMap project, which has harmonized cell lines and drug names^32,55^. In total 974 cell lines tested across 398 drugs are included in this dataset, amounting to 387,626 cell line-drug-IC_50_ pairs (CDI pairs). After intersecting for cell lines included in the CCLE RNA-seq compendium, selecting drugs for which we could obtain SMILES string, removing CDI pairs representing combination therapies and pairs with missing values for either drug name or IC_50_, 692 cell lines tested on 310 drugs remained, amounting to 185,186 CDI pairs. All IC_50_ values were transformed to log scale log_10_(IC_50_). After selecting for cell lines that represent the same tissue of origin as the TCGA dataset (25 tumor types), 531 cell lines tested on 310 drugs amounting to 142,351 CDI pairs.

## GLOSSARY

*GCN*: Graph Convolutional Network
*MorganFP*: Morgan Fingerprint
*SMILES*: Simplified Molecular Input Line Entry System for annotating chemical structures using character strings
*ML/DL*: machine learning/deep learning
*CDR*: Clinician Drug Response
*CDI*: Cell-line-Drug-IC_50_
*EM*: Expression Module
*DM*: Drug Module
*PM*: Prediction Module
*OOD*: Out of Distribution
*CL*: Cell Line

## AUTHOR CONTRIBUTIONS

I.A. conceived the idea, performed deep learning framework modeling, optimization and analysis. L.S. contributed in developing ideas on how to align cell lines and patients. H.D., J.S. supervised the project. All authors prepared the manuscript.

## CODE AVAILABILITY

The package and API for PACE is available at https://github.com/ioannisa92/PACE. The code and data to train and deploy the model of this manuscript are available on the github.

